# A new class of cell wall-recycling L,D-carboxypeptidase determines β-lactam susceptibility and morphogenesis in *Acinetobacter baumannii*

**DOI:** 10.1101/2021.09.22.461454

**Authors:** Yunfei Dai, Victor Pinedo, Amy Y. Tang, Felipe Cava, Edward Geisinger

## Abstract

The hospital-acquired pathogen *Acinetobacter baumannii* possesses a complex cell envelope that is key to its multidrug resistance and virulence. The bacterium, however, lacks many canonical enzymes that build the envelope in model organisms. Instead, *A. baumannii* contains a number of poorly annotated proteins that may allow alternative mechanisms of envelope biogenesis. We demonstrated previously that one of these unusual proteins, ElsL, is required for cell elongation and for withstanding antibiotics that attack the septal cell wall. Curiously, ElsL is composed of a leaderless YkuD-family domain usually found in secreted, cell-wall-modifying L,D-transpeptidases (LDTs). Here, we show that, rather than being an LDT, ElsL is actually a new class of cytoplasmic L,D-carboxypeptidase (LDC) that provides a critical step in cell-wall recycling previously thought to be missing from *A. baumannii*. Absence of ElsL impairs cell wall integrity, elongation, and intrinsic resistance due to buildup of murein tetrapeptide precursors, toxicity of which is bypassed by preventing muropeptide recycling. Multiple pathways in the cell become sites of vulnerability when ElsL is inactivated, including L,D-crosslink formation, cell division, and outer membrane lipid homoeostasis, reflecting its pleiotropic influence on cell envelope physiology. We thus reveal a novel class of cell-wall-recycling LDC critical to growth and homeostasis of *A. baumannii* and likely many other bacteria.

**Importance:** To grow efficiently, resist antibiotics, and control the immune response, bacteria recycle parts of their cell wall. A key step in the typical recycling pathway is the reuse of cell wall peptides by an enzyme known as an LDC. *Acinetobacter baumannii*, an “urgent-threat” pathogen causing drug-resistant sepsis in hospitals, was previously thought to lack this enzymatic activity due to absence of a known LDC homolog. Here, we show that *A. baumannii* possesses this activity in the form of an enzyme class not previously associated with cell wall recycling. Absence of this protein intoxicates and weakens the *A. baumannii* cell envelope in multiple ways due to the accumulation of dead-end intermediates. Several other organisms of importance to health and disease encode homologs of the *A. baumannii* enzyme. This work thus reveals an unappreciated mechanism of cell wall recycling, manipulation of which may contribute to enhanced treatments targeting the bacterial envelope.

## Introduction

The Gram-negative pathogen *Acinetobacter baumannii* is a significant cause of healthcare-associated infections, including ventilator-associated pneumonia, bloodstream infections, urinary tract infections, and sepsis (1, 2). *A. baumannii* strains show widespread multidrug-resistance, limiting the number of therapies effective against such infections (3). This problem is compounded by the insufficient pipeline of new antibiotics active toward Gram-negative pathogens (4). Reflecting the urgency of this threat, the World Health Organization has ranked *A. baumannii* as a pathogen of highest priority for research and development of new antibiotics (4).

The distinct cell envelope of *A. baumannii* is a key potential target for new treatments, but many aspects of its synthesis and control are not understood. Building the critical peptidoglycan (PG) cell wall layer in particular appears to involve several unconventional and poorly defined strategies. A number of proteins integral to the canonical pathways of septal PG synthesis, cell separation, and PG recycling have no homologs in the microorganism (5). For instance, *A. baumannii* lacks an ortholog of *E. coli* LdcA or *V. cholerae* LdcV, L,D-carboxypeptidases (LDCs) necessary for reusing peptides derived from old PG for synthesis of new cell wall (6, 7).

In addition to lacking canonical enzymes, the pathogen encodes several proteins that are implicated in PG homeostasis based on domain annotations but whose actual functions remain mysterious (5). Prominent among this group are two YkuD-domain proteins, which we have named ElsL (ACX60_RS03475) and Ldt_Ab_ (ACX60_RS05685) (5) (referred to as LdtK and LdtJ in (8)). YkuD domains are found in secreted enzymes that modify the cell wall for a variety of purposes in other organisms (9). For example, many YkuD-domain proteins carry out L,D-transpeptidase (LDT) reactions that generate 3-3 cell wall cross-links. These bonds are usually less abundant than the 4-3 cross-links catalyzed by DD-transpeptidases (PBPs) but may reinforce the wall against envelope stress (9). Unlike PBPs, LDTs are generally insensitive to β-lactam antibiotics (with the exception of carbapenems) and thus are potential sites of drug resistance (10, 11). Ldt_Ab_, but not ElsL, was recently found to be essential for 3-3 cross-links (8). Interestingly, ElsL does not contain a detectable secretion signal or additional domains such as PG binding motifs that would target the protein to the cell wall (5). Functional predictions based on identification of conserved domains in ElsL are therefore limited. A few leads were obtained based on genome-wide profiling of antibiotic susceptibility phenotypes (5). These studies found that ElsL deficiency is closely related to malfunction of the Rod system, the multiprotein PG synthetic machinery responsible for cell elongation. Mutations affecting ElsL and the Rod system both caused hypersensitivity to the same subset of β-lactam antibiotics as well as dramatic loss of the bacterium’s characteristic short-rod shape (5). ElsL mutation was also associated with increased outer membrane (OM) shedding (8). The role ElsL plays in the cell and its link to Rod system function and the OM remain unknown.

In this paper, we have determined the function of ElsL in *A. baumannii* cell wall synthesis. We show that ElsL defines a novel, noncanonical class of cytoplasmic LDC essential to cell wall recycling. We have identified critical perturbations to cell wall metabolism and integrity that occur in the absence of this function. We also delineated the complete network of genetic interactions with *elsL*, revealing multiple cellular pathways that are affected by its inactivation.

## Results

### ElsL and Ldt_Ab_ have opposing effects on L,D-crosslink formation in *A. baumannii*

To determine the effect of the two *A. baumannii* YkuD-family proteins, ElsL and Ldt_Ab_, on modifying the cell wall, we analyzed deletion mutants using two assays that detect PG remodeling: (1) metabolic cell wall labelling using the fluorescent precursor HCC-amino D-alanine (HADA), incorporation of which depends on an exchange reaction catalyzed by transpeptidases including LDTs (12); and (2) analysis of cell wall muropeptide composition by ultra-performance liquid chromatography/mass spectrometry (UPLC-MS) (13).

WT bacteria incubated with HADA incorporated the label throughout their cell walls, with the population of dividing cells showing increased signal at mid-cell (Fig. 1A, B). Deletion of *ldt*_Ab_ resulted in a dramatic loss of signal along the side-wall of cells that was reversed by reintroducing the gene *in trans* (Fig. 1A-D). By contrast, deletion of *elsL*, while causing loss of short rod shape, caused no decrease in HADA incorporation (Fig. 1A, B). These phenotypes confirm previous findings with a related fluorescent label (8). The Δ*ldt*_Ab_ phenotype also resembled that seen with *E. coli* completely lacking its 6 LDT paralogs (14).

**Fig. 1.**
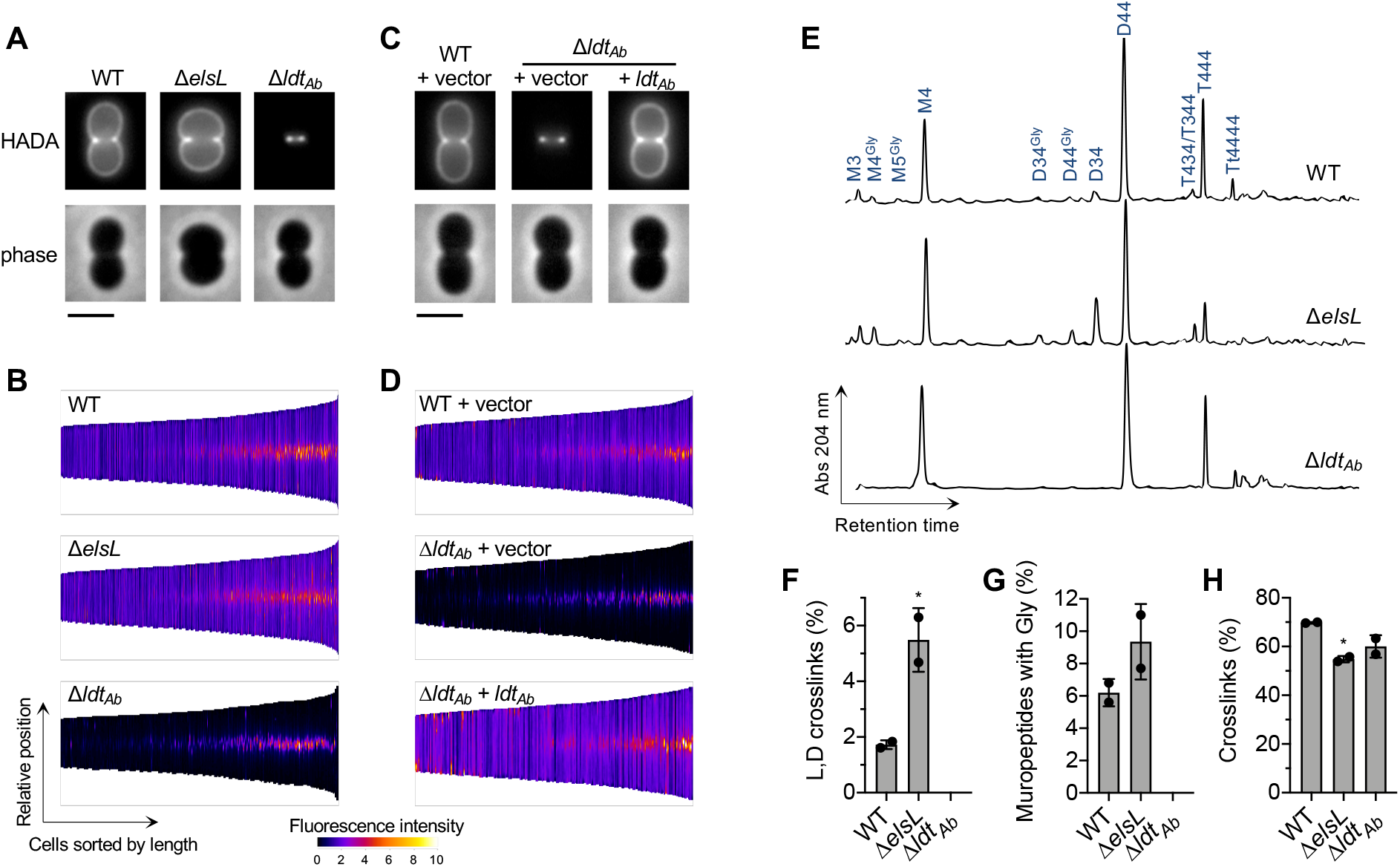
ElsL and Ldt_Ab_ have opposing effects on L,D-crosslink formation. (A and C) Cells of the indicated *A. baumannii* strain were metabolically labelled with HADA and imaged by fluorescence and phase-contrast microscopy. Representative cells are shown. Scale bar, 2µm. (B and D) Demographic representation of the cellular distribution and intensity of the HADA label. Cells (n ≥ 227 with each strain) were ordered according to their length and stacked by aligning cell midpoints. Fluorescence intensity along the medial axis of each cell is displayed as heat map. (E) Muropeptide profile analysis. Major muropeptides and those showing differences between WT and mutant are labelled. “M” indicates monomeric muropeptide; “D,” “T,” and “Tt” indicate dimeric, trimeric, or tetrameric cross-linked muropeptides, respectively; number(s) indicate the residue length of the stem peptide(s). “Gly” indicates that the terminal residue is a glycine. (F-H) Percentage of L,D-crosslinks, glycine-containing muropeptides, and total crosslinked muropeptides were quantified. Bars show mean ± s.d. (n = 2 biological replicates). *, p < 0.05 in unpaired t-test comparing Δ*elsL* vs WT (F-G) or in one-way ANOVA with Dunnett’s multiple comparisons test comparing each mutant to WT (H).

Cell wall muropeptide profiling revealed changes consistent with the metabolic labeling experiments. The WT profile consisted of major and minor muropeptide peaks similar to those reported previously (Fig. 1E) (15-18). Δ*ldt*_*Ab*_ caused complete loss of LDT-generated muropeptides, including the D34 (L,D-crosslinked) dimer (Fig. 1E, F) and muropeptides containing a terminal glycine (fourth position) residue from D-alanine exchange (Fig. 1E, G), consistent with prior findings (8). By stark contrast, the Δ*elsL* strain showed an increase in the L,D-crosslinked D34 muropeptide compared to WT (Fig. 1E, F; Fig. S1). Muropeptides with terminal glycines also appeared to increase in the Δ*elsL* mutant (Fig. 1E), although their overall levels were not significantly different from WT based on unpaired t-test (Fig. 1G). Despite the increase in 3-3-crosslinks, the Δ*elsL* cell wall showed lower total cross-linkage (including 4-3 as well as 3-3 bonds, Fig. 1H). Taken together, these data confirm that Ldt_Ab_ is the LDT responsible for alternative crosslinking within the *A. baumannii* cell wall, and indicate that ElsL is not a canonical LDT. Rather, ElsL has an alternative function that influences both 4-3 and 3-3 crosslink formation as well as cell shape.

Analysis of fluorescent protein fusions supports the prediction that ElsL function is cytosolic. Fusion of ElsL to a monomeric superfolder GFP (msGFP2 (19)) caused a diffuse signal throughout the cell interior, consistent with cytoplasmic localization (Fig. 2A). An analogous Ldt_Ab_-msGFP2 fusion, by contrast, showed peripheral fluorescent patches consistent with its predicted localization in the periplasm (Fig. 2B). This difference in localization occurred despite both gene fusions resulting in similarly efficient levels of intact chimeric proteins (Fig S2A). Expression of *elsL-msGFP2* reversed the shape defect of Δ*elsL* bacteria, resulting in short rods with maximal width matching that of WT, indicating the fusion protein was functional (Fig. 2C). With a cytoplasmic location partitioned away from the sacculus, ElsL may thus act as an LDC, an alternative or additional activity seen with some YkuD-family proteins (20, 21). The logical extension of this prediction is that ElsL is the missing-link cytoplasmic LDC allowing recycling of imported cell-wall fragments in *A. baumannii*.

**Fig. 2.**
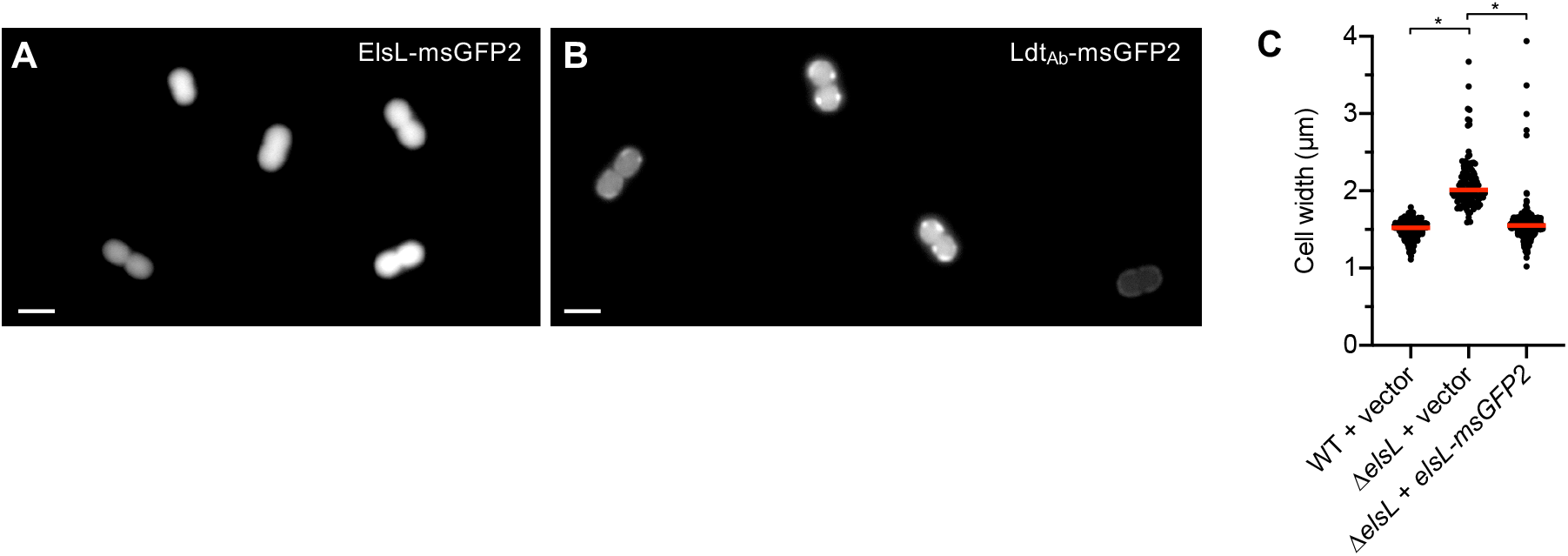
An ElsL-msGFP fusion shows diffuse cytoplasmic localization, while Ldt_Ab_-msGFP localizes to the cell periphery. (A-B) Fluorescent micrographs showing representative WT cells harboring the plasmid-borne gene fusions of msGFP2 to *elsL* (A) or *ldt*_*Ab*_ (B). Scale bar, 2 µm. (C) The *elsL-msGFP2* fusion reverses the morphology defect of Δ*elsL*. Cells were imaged by phase-contrast microscopy and cell width measured by image analysis. Bars show median values (n ≥ 118). *, p <0.0001 in Kruskal-Wallis test.

### *elsL* and *ldt*_*Ab*_ have interconnected aggravating genetic interactions

As a parallel approach to identify leads on ElsL function, we mapped its full network of genetic interactions throughout the genome, which ultimately enabled a series of tests of the above hypothesis. To this end, we performed comparative Tn-seq analysis using ElsL^+^ and ElsL^-^ strains. A dense *mariner* transposon library was constructed in the Δ*elsL* background, and colony growth of the resulting double mutants was quantified in massively parallel fashion by measuring Illumina sequencing read abundance corresponding to every member of the library (Materials and Methods). This growth was then compared to that of the matching single (ElsL^+^) transposon mutants within a control *mariner* library generated previously in the WT (5). Potential aggravating interactions were identified as genes for which knockout showed growth dependence on ElsL (i.e., low ratio of Δ*elsL*/control read counts), with significance determined by permutation test. Examination of such interactions should illuminate the pathways in which ElsL functions.

A large number of genes showed potential aggravating interactions with *elsL* (Fig. 3A, left; Dataset S1). Several of these mapped to cell wall synthesis, cell division, and envelope stability pathways. For example, the non-essential cell division-associated genes *minCD, blhA* (5, 22) and ACX60_RS13190 (structural maintenance of chromosomes family (5)), showed substantially lower Tn-seq read counts when mutated in combination with Δ*elsL* (Fig. 3A, B). A similar result was seen with *zapA*, a cell division locus linked to *RS13190*, albeit without passing the 5% false-discovery rate cut-off (Fig. 3A, B). In addition, nearly all *mla* genes, which determine lipid transport in the OM (23, 24), had Tn-seq read counts dependent on intact *elsL* (Fig. 3A,B). Notably, *ldt*_*Ab*_ also showed a prominent growth defect dependent on Δ*elsL* (Fig. 3A,B), indicating that knockout of both YkuD-family proteins was poorly tolerated in *A. baumannii*, as suggested previously (8). To interrogate this last result, we generated a parallel *mariner* library in the Δ*ldt*_*Ab*_ strain and analyzed its genome-wide interactions (Dataset S2). While many fewer potential negative interactions were identified with Δ*ldt*_*Ab*_ compared to those with Δ*elsL*, the strongest hit was *elsL* (Fig. 3A, right; Fig. 3B; Dataset S2), supporting the notion that the two genes have an aggravating interaction.

**Fig. 3.**
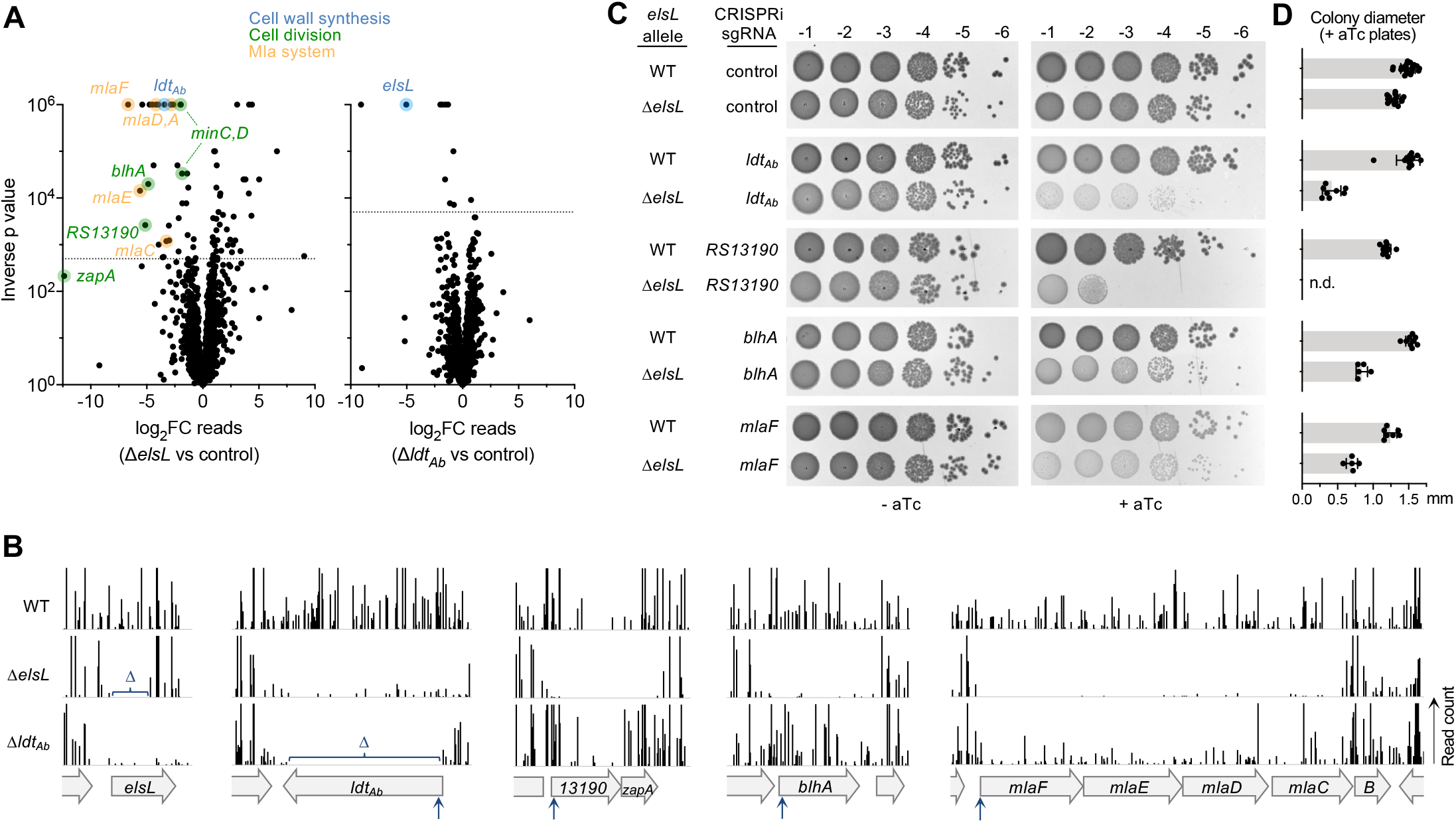
Tn-seq reveals aggravating genetic interactions with *elsL* and *ldt*_Ab_. (A) Tn-seq genetic interaction analysis. Volcano plot shows the ratio of Tn-seq read counts mapped to genes in the mutant *mariner* transposon library (Δ*elsL or* Δ*ldt*_*Ab*_) compared to the control transposon library (WT). Dotted horizontal lines indicate a false discovery rate (q-value) of 0.05. Hits described in the text are color-highlighted according to the indicated pathway. (B) Tracks show Tn-seq read counts at each insertion site at different chromosomal loci within the indicated *mariner* library. Bars represent normalized read count and vertical arrows indicate position targeted by CRISPRi. Δ indicates a deleted region. (C) Validation of aggravating genetic interactions with Δ*elsL* via targeted CRISPRi and colony formation tests. WT or Δ*elsL* strains harboring aTc-inducible *dcas9* and the indicated sgRNA were serially diluted (10-fold) and spotted on LB agar medium supplemented with 0 or 50 ng/ml aTc. Colonies resulting after overnight 37°C incubation were imaged. (D) Diameters of colonies from triplicate aTc plates at 10^−6^ or higher dilution were measured by image analysis. Bars show mean ± s.d. (n ≥ 5 colonies). p < 0.0001 in unpaired t-tests comparing Δ*elsL* to WT with each sgRNA. n.d., colonies not detected and statistical test not performed.

We used CRISPRi (25) to validate the negative genetic interactions between *elsL* and the strongest hits within each pathway (*ldt*_*Ab*_, *RS13190-zapA, blhA*, and *mlaC-F*). Chimeric single guide RNAs (sgRNAs) were designed to target the 5’ end of each locus (Fig. 3B, blue arrows) and were introduced into WT and Δ*elsL* strains harboring anhydrotetracycline (aTc)-inducible dCas9. CRISPRi allows efficient knockdown of operons (26); *RS13190-zapA* and *mlaC-F* are therefore each likely to be co-repressed in this strategy. The effect of knockdown on colony growth was compared with two parallel controls—(1) growth in the absence of dCas9 inducer (-aTc), and (2) growth with a control, non-targeting sgRNA (25). CRISPRi of each locus in the WT strain resulted in colony growth that was at or near control levels (Fig. 3C, D). By contrast, CRISPRi in Δ*elsL* caused greatly amplified growth defects. Silencing *RS13190* in the Δ*elsL* background completely blocked colony formation (Fig. 3C, D), indicative of strong genetic aggravation. Other knockdowns in Δ*elsL* resulted in significantly reduced colony growth compared to controls (Fig. 3C, D). To test whether these growth defects reflect aggravating interactions with *elsL*, we compared them to those expected from a null multiplicative model based on the growth of single-lesion strains (Materials and Methods). Each double-lesion strain (CRISPRi + Δ*elsL*) had significantly lower growth than expected from the null model (Table S1). Together, these results indicate that *elsL* interacts negatively with LDT, cell division, and OM lipid transport pathways.

Aggravating interactions frequently arise if the underlying genes function in parallel or redundant pathways (27). This scenario could explain the basis for the aggravating interaction between *elsL* and nonessential cell division genes. Mutation of *blhA* (22) and silencing of *RS13190*-*zapA* (Fig. S3A) each impair cell division and cause a mode of growth that is more heavily dependent on cell elongation compared to WT. Loss of *elsL* blocks cell elongation (5) (Fig. 2C). Blocking genes of each type in combination thus likely results in cells unable to divide or elongate efficiently. This is supported by microscopy of Δ*elsL* CRISPRi_*RS13190*-*zapA*_, which showed large, irregular spheroid morphology rather than the filaments seen with CRISPRi_*RS13190*-*zapA*_ alone (Fig. S3A). These data are consistent with synthetic lethality occurring due to an inability of cells to continue growing their cell wall laterally as a way to compensate for deficient or delayed cell division. The defect in elongational growth machinery may also affect how septal PG synthesis is initiated (28, 29), hypersensitizing to inhibition of a parallel cell division pathway. In addition to parallel pathways, aggravating interactions may arise when one gene product limits the toxicity generated by absence of the other (30-32). We considered this alternative model in examining the mechanism of the *elsL*-*ldt*_*Ab*_ genetic interaction.

### Synthetic lethal relationship between *elsL* and *ldt*_*Ab*_

To investigate the *elsL*-*ldt*_*Ab*_ genetic relationship, we combined the two deletions (Materials and Methods) and identified conditions that further aggravate its growth phenotype. The Δ*elsL* Δ*ldt*_*Ab*_ double mutant formed small, translucent colonies, identical to the phenotype of the Δ*elsL* CRISPRi_*ldt*_*Ab* strain (Fig. 4A). Since both genes affect the cell wall, we examined growth with low osmolarity medium (LB without NaCl, “LB_0_”). LB_0_ medium dramatically aggravated the Δ*elsL* Δ*ldt*_*Ab*_ growth defect, with colony formation completely blocked at dilutions beyond 10^−2^, indicating synthetic lethality (Fig. 4A). A similar result was obtained with Δ*elsL* CRISPRi_*ldt*_*Ab* (Fig. S3B). Consistent with the colony phenotypes, Δ*elsL* Δ*ldt*_*Ab*_ showed lower viable counts during liquid culture compared to the single mutants and WT, with low osmolarity amplifying the defect (Fig. 4B). The Δ*elsL* Δ*ldt*_*Ab*_ sacculus showed compositional defects that reflected the combination of each single lesion, with low overall cross-linking and absence of LDT-mediated muropeptides (Fig. S3C, D). The detrimental effect of lacking both genes also manifested in defective cell shape, with the double mutant forming enlarged spheroids in LB and irregular, bloated shapes with frequent blebs and lysis in LB_0_ (Fig. 4C). Notably, the Δ*elsL* single mutant also had viability and shape defects compared to WT that were exacerbated by low osmolarity, while Δ*ldt*_*Ab*_ was unaffected (Fig. 4B,C), underlining the importance of ElsL to physiology and stress resistance. Together, these results indicate that *elsL* and *ldt*_*Ab*_ have a synthetic lethal relationship, with the *elsL* defect in particular making cells sensitive to conditions of increased cell wall stress.

**Fig. 4.**
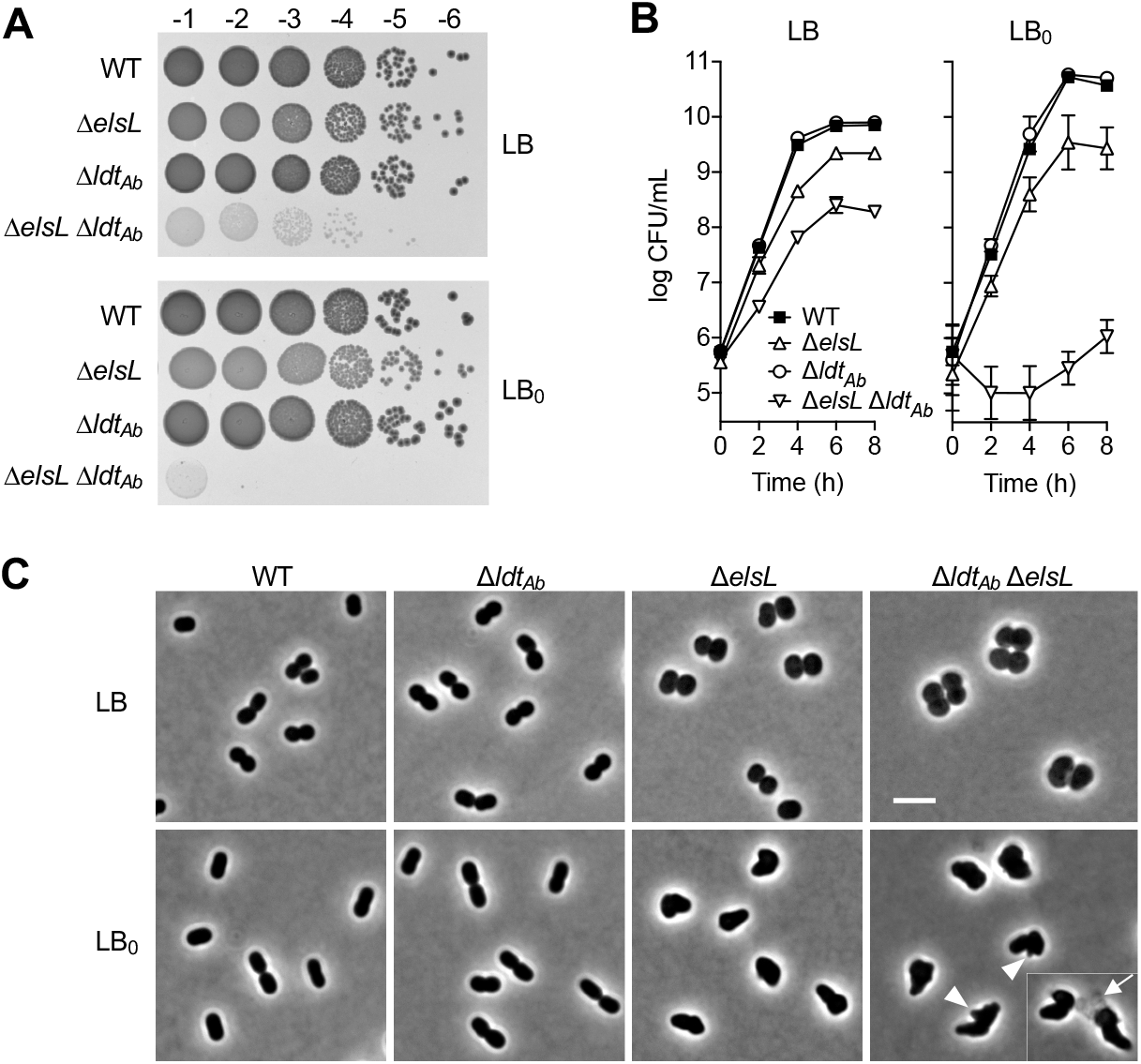
Synthetic lethal interaction between *elsL* and *ldt*_Ab_. (A) WT *A. baumannii* or the indicated deletion mutant were serially diluted and spotted on LB (top) or LB_0_ (bottom) agar medium, and the resulting colonies were imaged as in Fig. 3. (B) The indicated strains were cultured to saturation in LB, diluted in LB or LB_0_, and viable counts determined over time. Data points show geometric mean ± s.d. (n = 3 biological replicates). Where not visible, error bars are within the confines of the symbol. (C) The indicated strains (columns) cultured in LB or LB_0_ (rows) were imaged by phase-contrast microscopy. Arrowheads indicate blebs; arrow indicates example of lysed cell lacking dense phase contrast. Scale bar, 4 µm.

### Suppression of *elsL*-*ldt*_*Ab*_ synthetic lethality by blocking cell wall muropeptide recycling

To illuminate ElsL function and identify the source of Δ*elsL* toxicity, we exploited the synthetic lethality of Δ*elsL* Δ*ldt*_*Ab*_ and isolated suppressor mutants reversing its major growth defect. Large, opaque colonies forming from Δ*elsL* Δ*ldt*_*Ab*_ on solid LB or LB_0_ medium were purified and their mutations mapped (Materials and Methods). Of 22 distinct suppressors identified using this strategy, 21 mapped to 2 genes functioning in PG recycling, *ampG* (the muropeptide permease) and *mpl* (murein peptide-UDP-MurNAc ligase) (33) (Fig. 5A). In each case, the mutation was a predicted null allele. Representative mutants showed enhanced colony growth with both LB and LB_0_ (Fig. 5B), as did an independently constructed, in-frame deletion of *ampG* (Fig. 5C). The remaining suppressor, which also enhanced colony growth relative to its parent (Fig. 5B), mapped to ACX60_RS13100, which we have named *ltgF* (lytic trans<u>g</u>lycosylase determining <u>f</u>osfomycin susceptibility, as explained below). This locus encodes a predicted lytic transglycosylase having an MltD-like catalytic domain and a large C-terminal region with multiple LysM repeats resembling the PG-anchoring domain of autolysins (Fig. 5D) (34). The mutant allele had an insertion sequence (IS) between these two regions. As a predicted PG turnover enzyme, *ltgF* may have an important role in PG recycling like the other sites of suppression. Supporting this idea, the Tn-seq antibiotic susceptibility profile (phenotypic signature) (5) of *ltgF* closely correlates with those of canonical PG recycling genes (Fig. 5E, Fig. S4A). In-frame deletion of *ltgF* in ATCC 17978 (Fig. 5F, G) as well as transposon mutation in a different strain background (AB5075, Fig. S4B, C), each caused hypsersusceptibility to Fosfomycin, a mark of defective cell wall recycling (35, 36). The susceptibility defect was akin to that seen with recycling-blocked *ampG* mutants tested in parallel (Fig. 5F, G; Fig. S4B, C). These results support a role for *ltgF* in PG turnover and recycling. The findings altogether reveal that the Δ*elsL* Δ*ldt*_*Ab*_ growth defect is suppressed by blocking cell wall recycling at key points— its earliest steps (generation of anhydromuropeptides or their import) or at the step of peptide reuse (Fig. 5I, left, bold steps).

**Fig. 5.**
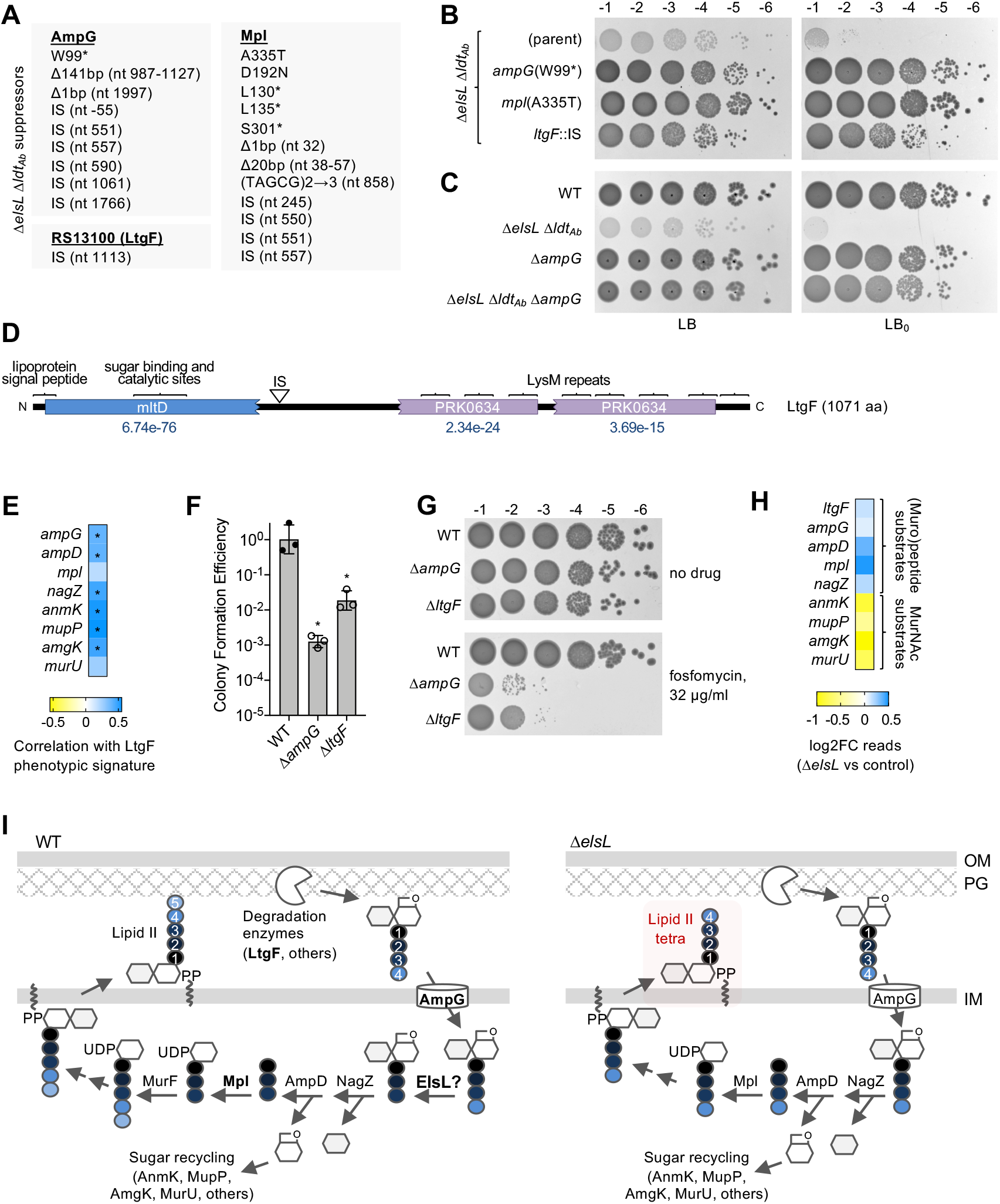
Blocking early cell wall recycling proteins or the Mpl peptide-recycling ligase suppresses Δ*elsL* Δ*ldt*_*Ab*_ synthetic lethality. (A) Listed are mutations identified by whole-genome sequencing in derivatives of Δ*elsL* Δ*ldt*_*Ab*_ allowing enhanced colony formation on LB or LB_0_ agar medium. Protein effect is listed in the case of substitutions; in all other cases, the mutation and the corresponding nt position relative to the gene start are listed. IS, insertion sequence. (B-C) The indicated spontaneous mutant and the Δ*elsL* Δ*ldt*_*Ab*_ parent strain (B), or the indicated deletion mutants and WT control (C) were serially diluted and spotted on LB (left) or LB_0_ (right) agar medium, and the resulting colonies imaged as in Fig. 3. (D) Schematic of LtgF protein. IS indicates location affected by the ISAba1 suppressor mutation in *ltgF*. Rectangles and brackets indicate predicted domains from CDD or SignalP. CDD e-values are listed; e-values of the LysM domains were below 1e-5. (E) LtgF and canonical cell wall recycling genes have correlated phenotypic signatures. Heat map shows Pearson correlation coefficient (r) measuring relatedness of the Tn-seq phenotypic signatures (5) of each gene with that of *ltgF*. *, p < 0.05. (F-G). Susceptibility to fosfomycin (32 µg/ml) was determined by CFE assay. Bars (F) show geometric mean ± s.d (n = 3 biological replicates). *, p < 0.0001 in unpaired t-tests comparing each mutant to WT. Representative colonies are shown in G. (H) Direction of the Tn-seq genetic interaction between *elsL* and cell wall recycling genes depends on the component being recycled. Heat map shows fold change in Tn-seq read counts of the indicated gene in the Δ*elsL mariner* library vs in the WT control library (Dataset S1). (I) Model for role of ElsL in *A. baumannii* cell wall recycling. (Left) PG is degraded by lytic transglycosylases (e.g., LtgF), and endopeptidases to release anhydro-MurNAc-containing products. Anhydro-MurNAc-tetrapeptides imported by AmpG are processed to tripeptide form by ElsL. After removal of the PG amino sugars, the tripeptide is reused via ligation to UDP-MurNAc by Mpl, followed by addition of D-Ala-D-Ala to result in UDP-MurNAc-pentapeptide and ultimately lipid II. In the absence of ElsL, by contrast, tetrapeptides are reused by Mpl, resulting in synthesis of aberrant lipid II tetrapeptide (Right).

Re-examination of our genetic interaction Tn-seq data revealed that blocking the same cell wall recycling steps identified as suppressors of Δ*elsL* Δ*ldt*_*Ab*_ lethality also alleviated the mild colony growth defect of Δ*elsL*. For instance, transposon mutations in *ltgF* and cell wall recycling enzymes acting on muropeptide or peptide substrates (Fig. 5I) each led to slightly enhanced growth within the Δ*elsL* library compared to control (Fig. 5H). By contrast, mutation of other recycling steps dedicated to MurNAc sugar recycling had the opposite effect, decreasing Δ*elsL* growth compared to control (Fig. 5H). A clear differential effect of blocking different branches of the cell wall recycling circuit on growth was not observed with the Δ*ldt*_*Ab*_ library (Fig. S4D). These results point to ElsL deficiency and peptide reuse as the key sources of synthetic toxicity that are bypassed by cell wall recycling block.

The suppressor analysis clearly implicates ElsL in cell wall recycling. Integrated with our genetic interaction, muropeptide composition, and localization data, these results lend strong support for the above model that ElsL is the cell wall recycling LDC in *A. baumannii*. The deleterious effects of ElsL deficiency can thus be explained as a consequence of aberrant buildup of its tetrapeptide substrates. Processing of these intermediates by Mpl and other enzymes would lead to generation of lipid-II tetrapeptide and ultimately incorporation of tetrapeptide stems into the cell wall (Fig. 5I, right). These stems are toxic dead-ends, as they cannot be used as donors in cross-link formation by PBPs (37). This would account for the decrease in overall crosslinks seen with Δ*elsL* (Fig. 1E, H). Toxicity would be limited to some extent by Ldt_Ab_, since LDTs use tetrapeptides as donors (6); this would account for the elevated level of L,D-crosslinks seen with Δ*elsL* (Fig. 1F). Absent Ldt_Ab_, however, the uncrosslinked tetrapeptide stems would compromise cell wall integrity, accounting for the aggravated phenotypes (Fig. 4). Preventing tetrapeptide reuse by suppressor mutations would efficiently bypass this toxic pathway.

### Block in cell wall recycling suppresses Δ*elsL* shape and susceptibility defects

We interrogated the ElsL LDC model through parallel genetic and biochemical approaches. First, given the ability of muropeptide recycling block to enhance Δ*elsL* Tn-seq fitness (Fig. 5H), we tested the degree to which such block also alleviates Δ*elsL* morphological and antibiotic hypersusceptibility phenotypes. Blocking cell wall recycling by *ampG* mutation completely restored a short rod shape to cells lacking ElsL, with median cell width returning to the WT value (Fig. 6A, B). Δ*elsL* bacteria are hypersusceptible to antibiotics that attack division septum PG synthesis, such as sulbactam (5). *ampG* deletion also completely reversed this defect. While Δ*elsL* was unable to form colonies on solid medium with low doses of sulbactam, Δ*elsL* Δ*ampG* grew efficiently, restoring the MIC to a WT level (1µg/ml; Fig. 6C). This effect extended to another treatment predicted to target septal PG synthesis, the broad-spectrum combination drug piperacillin-tazobactam (38). Specific block of divisional PG synthesis by piperacillin-tazobactam was supported by the dramatic elongation of WT cells in the presence of the drug at sub-MIC, and by the marked hypersusceptibility of an elongation-defective Rod system mutant, Δ*pbp2* (Fig. S5A, B). Δ*elsL* similarly showed increased susceptibility to piperacillin-tazobactam (Fig. S5B). Hypersusceptibility was fully reversed by reintroducing a WT *elsL* allele (Fig. S5C) or, notably, by removing cell wall recycling through deletion of *ampG* (Fig. 6D). Together, these results are consistent with ElsL deficiency owing its multiple toxic phenotypes to recycling of aberrant cell wall intermediates.

**Fig. 6.**
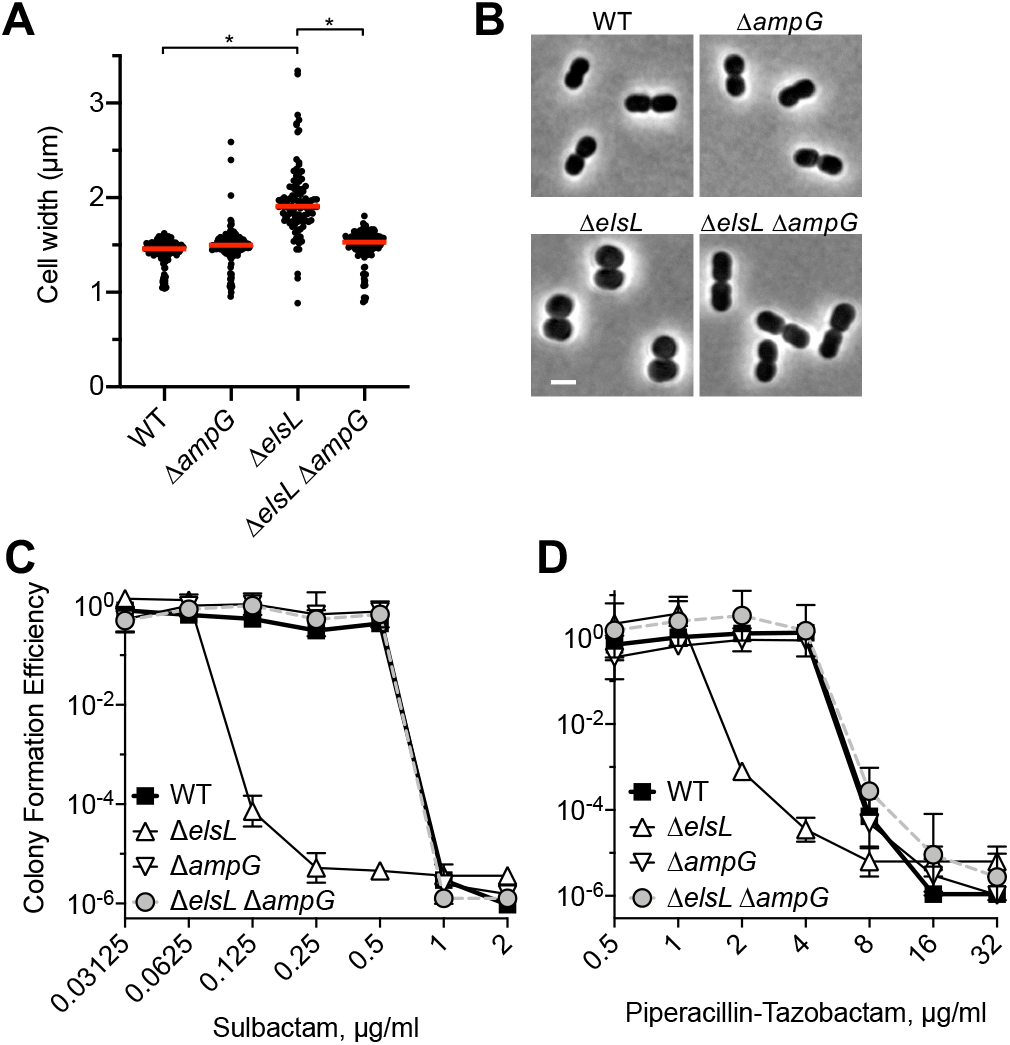
Blocking cell wall recycling suppresses morphology and antibiotic hypersusceptibility defects of Δ*elsL* mutant. (A-B) *ampG* deletion restores rod shape in the Δ*elsL* strain. Cells were imaged by phase-contrast microscopy and cell width measured by image analysis (A). Lines show median values (n ≥ 102). *, p <0.0001 in Kruskal-Wallis test. Representative cells are shown in B. Scale bar, 2 µm. (C-D) *ampG* deletion reverses Δ*elsL* hypersusceptibility to divisome-targeting β-lactams. Susceptibility to sulbactam (C) and piperacillin-tazobactam (D) was measured by CFE assay. Data points show geometric mean ± s.d. (n = 3 biological replicates).

### ElsL deficiency causes accumulation of aberrant cytosolic tetrapeptides and disrupts the synthesis of pentapeptide precursors

We further tested the potential recycling LDC function of ElsL by measuring its effect on levels of PG precursors *in vivo*. Canonical recycling LDCs act on tetrapeptide-containing products derived from turnover of the cell wall (6, 7). The resulting tripeptides lead to generation of the key cytosolic building block UDP-MurNAc pentapeptide (UDP-M5) (Fig. 5I, left); aberrant reuse of unprocessed tetrapeptides, by contrast, would lead to the potentially toxic UDP-MurNAc tetrapeptide (UDP-M4) (Fig. 5I, right). UDP-M4 should therefore show higher levels in ElsL^-^ vs ElsL^+^ *A. baumannii* if the protein functions as an LDC. To test this prediction, we analyzed UDP-linked muramyl-peptides within the bacterial strains via UPLC-MS. In contrast to ElsL^+^ strains, which had UDP-M5 without detectable UDP-M4 (Fig. 7A, WT and Δ*ldt*_*Ab*_), ElsL^-^ mutants showed abundant UDP-M4 with a concomitant decrease in UDP-M5 (Fig. 7A, Δ*elsL* and Δ*elsL* Δ*ldt*_*Ab*_), consistent with tetrapeptide substrate buildup due to an absence of LDC function. Analysis of isogenic AmpG^-^ strains revealed the likely mechanism of suppression of blocking cell wall recycling. Compared to AmpG^+^, AmpG^-^ variants showed dramatically reduced amounts of UDP-M4 that were now below the level of UDP-M5 (from *de novo* biosynthesis) in each strain (Fig 7A, Δ*elsL* Δ*ampG* and Δ*elsL* Δ*ldt*_*Ab*_ Δ*ampG*). AmpG deletion therefore prevents the toxic tetrapeptide products from being reused for PG synthesis, explaining how recycling mutations bypass the toxic phenotypes of ElsL deficiency.

**Fig. 7.**
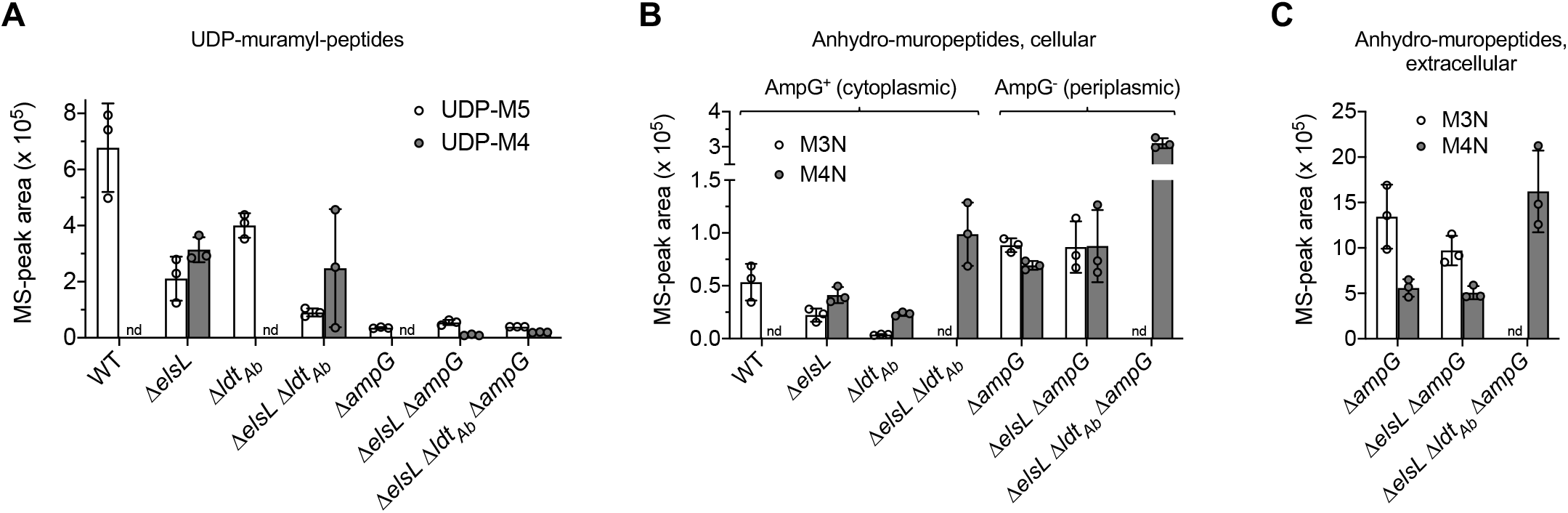
ElsL absence causes accumulation of cytoplasmic tetrapeptide recycling intermediates and a reduction in pentapeptide precursor synthesis that is bypassed by *ampG* deletion. (A) UDP-muramyl-peptides (UDP-M5, UDP-M4) were detected by MS in the cellular fraction from cultures of the indicated strains. (B-C) Anhydro-muropeptides (M3N and M4N) were detected in the intracellular (B) and extracellular (supernatant, C) fractions by MS from cultures of the indicated strains. In panel B, anhydro-muropeptides detected in AmpG^+^ cultures are imported and thus reflect cytoplasmic levels; detection of these PG turnover products in AmpG^-^ cultures are not efficiently imported and thus reflect levels in the periplasm. Bars show mean ± s.d (n = 3 biological replicates). nd, not detected.

Examination of anhydromuropeptides, the turnover-derived fragments that are recycled into UDP-linked precursors, further supports the critical function of ElsL as a recycling LDC. In WT, anhydromurotripeptide (M3N) but not anhydromurotetrapeptide (M4N) was detected in the cellular fraction (Fig. 7B). By contrast, Δ*elsL* had a buildup of M4N (Fig. 7B), consistent with lack of a recycling LDC. The AmpG permease is required for cytosolic import of anhydromuropeptides (33); detection of these products in AmpG^-^ cultures thus reflects a periplasmic or extracellular, rather than cytoplasmic, locale. By contrast to the results with AmpG^+^ strains, *elsL* deletion did not affect periplasmic (Fig. 7B) or extracellular (Fig. 7C) levels of M4N in the Δ*ampG* background (compare Δ*elsL* Δ*ampG* to Δ*ampG*). These results are consistent with ElsL providing only a cytoplasmic LDC function. In addition to these findings, we detected some M3N in Δ*elsL*, as well as an M4N increase/M3N decrease vs WT with Δ*ldt*_*Ab*_ (Fig. 7B). In each case, the levels of these anhydromuropeptides are largely a reflection of the compositionally altered cell walls from which they derive (e.g., the Δ*ldt*_*Ab*_ wall lacks tripeptide stems while the Δ*elsL* accumulates them; Fig. 1E). In the case of Δ*elsL* Δ*ldt*_*Ab*_, the large increase in M4N and undetectable M3N (Fig. 7B, C) are consistent with the combined effect of no cytosolic LDC activity and altered cell wall composition.

### ElsL has LDC activity paralleling that of *E. coli* LdcA

Three additional results support the model that ElsL is the missing LDC in *A. baumannii*. First, Δ*elsL* defects are reversed by LdcA, the canonical LDC from *E. coli*. To show this, each protein was 3X-FLAG-tagged and expressed in Δ*elsL* under IPTG control, and IPTG concentrations yielding equivalent protein levels were determined (50 µM with ElsL_FLAG_; 500 µM with LdcA_FLAG_; Fig. 8A; Fig. S2B). Using these conditions, both enzymes completely reversed the sulbactam hypersusceptibility of Δ*elsL* (Fig. 8B). At lower expression levels, LdcA caused partial complementation, while full complementation was still seen with ElsL (Fig. 8A, B; compare *ldcA*_FLAG_ 50 with *elsL*_FLAG_ 0 µM IPTG [leaky expression]). These results confirm that the Δ*elsL* defect is due to deficient LDC activity, and suggest that *A. baumannii* cell wall recycling has evolved to be most efficient with the ElsL YkuD-family LDC.

**Fig. 8.**
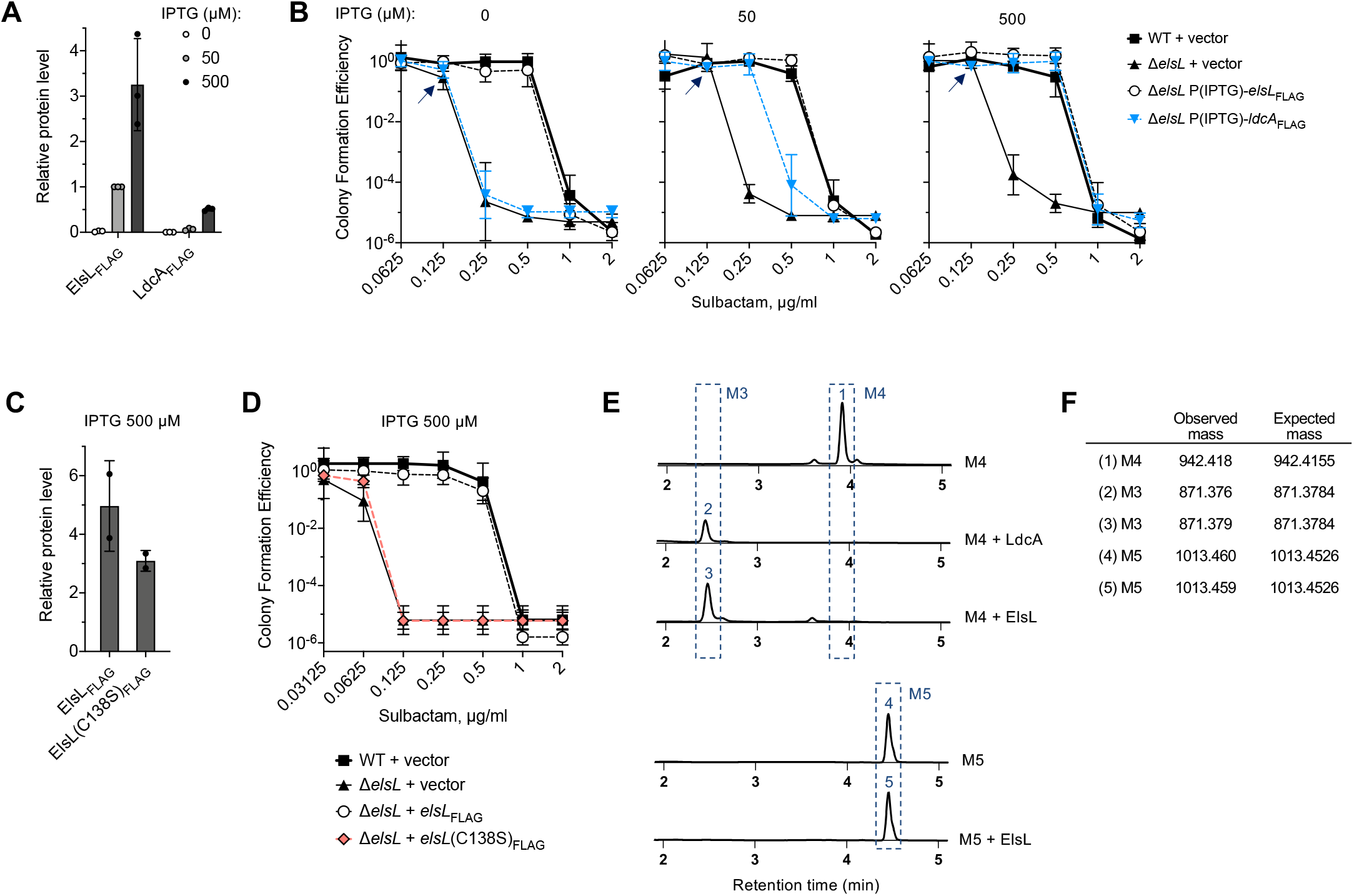
ElsL is the missing link PG recycling L,D-carboxypeptidase in *A. baumannii*. **(A)** Western blot quantification of ElsL_FLAG_ and LdcA_FLAG_. Δ*elsL* bacteria with inducible expression of each enzyme by P(IPTG) were cultured with the indicated IPTG concentration. Data points show protein level (relative to ElsL_FLAG_, IPTG 50 µM) determined from image analysis. Bars show mean ± s.d. (n = 3 biological replicates). (B) *E. coli* LdcA reverses the Δ*elsL* sulbactam hypersusceptibility defect. Susceptibility was determined by CFE assay. Plates contained the indicated sulbactam and IPTG concentrations. Data points show geometric mean ± s.d. (n = 3 biological replicates). Arrows denote that colonies formed by Δ*elsL* + vector on 0.125 µg/ml sulbactam medium were pinpoint-sized. (C) Western blot quantification of ElsL_FLAG_ in Δ*elsL* bacteria with the indicated P(IPTG)-inducible allele cultured in the presence of 500 µM IPTG. Data points show protein level (relative to ElsL_FLAG_, IPTG 50 µM) as in A except n = 2. (D) Intrinsic sulbactam resistance conferred by ElsL depends on an active site cysteine. Susceptibility to sulbactam was determined by CFE as in B. (E-F) ElsL has LDC activity *in vitro*. Shown are UPLC chromatograms of the reaction products following incubation of substrate (M4, murotetrapeptide, left; or M5, muropentapeptide, right) with the indicated purified enzyme (E). Peaks (numbered) were identified based on retention time and confirmed by MS (F).

Second, ElsL activity depends on a cysteine residue that aligns with the conserved catalytic cysteine of YkuD-family LDTs (39, 40). ElsL harboring a mutation at this site (C138S) lost all ability to confer intrinsic sulbactam resistance to Δ*elsL* when expressed *in trans* (Fig. 8D), despite high-level induction with 500 µM IPTG (Fig. 8C, Fig. S2C).

Third, purified ElsL has LDC activity *in vitro*. ElsL and *E. coli* LdcA, a positive control for LDC activity, were both purified using a C-terminal His-tag and assayed in parallel. LDCs cleave between the D-alanine and the meso-diaminopimelic acid only in disaccharide tetrapeptides (M4) and not in disaccharide pentapeptides (M5); we thus used these two muropeptides to assay LDC activity. Similar to LdcA, ElsL completely cleaved the M4 substrate to the disaccharide tripeptide (M3) (Fig. 8E, F). ElsL had no activity on M5 (Fig. 8E, F). ElsL thus possesses selective LDC activity. We conclude that ElsL is a novel type of cell-wall recycling enzyme using a cytoplasmic YkuD-family domain to catalyze the key LDC reaction.

### The ElsL LDC family includes orthologs in diverse bacteria

Search of a database of ∼14K representative bacterial genomes (Materials and Methods) revealed predicted cytoplasmic ElsL orthologs in a range of different organisms (Dataset S3). Orthologs were found mainly within the proteobacteria, with largest representation by diverse members of the gamma- and beta-proteobacteria classes (624 and 101, respectively, out of 788 total hits identified). This included the intracellular pathogens *Legionella pneumophila* and *Coxiella burnetii* (41). 5/21 gamma- and 2/6 beta-proteobacterial orders did not contain ElsL orthologs, suggesting that while widespread, the enzyme may have been lost or replaced by other LDCs in those cases. ElsL orthologs were also present in all orders of the *Verrucomicrobia*, including the human gut microbiota member *Akkermansia muciniphila*. In sum, a large number of diverse bacterial species appear to have evolved to use the cytoplasmic ElsL-class LDC for recycling of the cell wall.

## Discussion

We report here the identification of a novel class of cytoplasmic LDC that enables cell wall recycling in *A. baumannii*. This class uses a cysteine-containing catalytic domain usually found in LDT proteins. ElsL is the founding member of this family, and orthologs were identified in a range of other bacteria. ElsL deficiency causes a number of phenotypes: (1) loss of rod shape; (2) impaired growth; (3) hypersensitivity to septal cell wall-targeting antibiotics; and (4) cell wall structural defects, including increased 3-3 cross-links with reduced overall cross-linkage. Most of these phenotypes are aggravated by mutating the enzyme necessary for 3-3 cross-link formation (Ldt_Ab_) or by low osmolarity, while they are suppressed by blocking muropeptide recycling. Each of these features is explained by the model that ElsL is the LDC that processes turnover-derived muropeptides in *A. baumannii*, with toxic Δ*elsL* phenotypes caused by the deleterious tetrapeptides that accumulate absent this processing. Analysis of cell wall precursors supported this model and showed that tetrapeptide-containing biosynthetic intermediates increase in Δ*elsL* bacteria at the expense of the mature pentapeptide building blocks (Fig. 7). When incorporated as part of nascent cell wall, these tetrapeptides cannot be used as donors in 4-3 bond formation by PBPs and thus diminish overall cell wall cross-linkage (Fig. 1H). Periplasmic LDTs, however, can act on tetrapeptides, likely mitigating their toxic effects and explaining the synthetic lethality of the double *elsL ldt*_*Ab*_ mutant (Fig. 4). ElsL has LDC activity *in vitro*, catalyzing the removal of the terminal D-Ala from disaccharide tetrapeptides but not pentapeptides (Fig. 8E), solidifying its critical role in muropeptide recycling.

The phenotypes created by absence of ElsL are highly similar to those seen with Rod system defects, pointing to a tight connection between ElsL LDC function and cell wall elongation. The shared phenotypes include loss of rod morphology as well as highly correlated antibiotic susceptibility signatures defined by marked hypersensitivity to division-targeting β-lactams (5). Our genetic interaction data showed that the Δ*elsL* growth defect was aggravated by mutation of non-essential cell division genes (Fig. 3), which mirrors the effect of antibiotics blocking division (Fig. 6). Together these results support the model that the Rod system is defective when cells lack ElsL, resulting in cells that are heavily reliant on divisome PG synthesis and hypersusceptible to inhibition of this process. Given its diffuse cytoplasmic location, the recycling LDC is almost certainly not a requisite component of the Rod complex; instead, it is accumulation of tetrapeptide intermediates that is likely responsible for Rod system failure and the specific deficiency in elongation. That aberrant intermediates may have selective effects on a particular PG biosynthetic machine is supported by previous work with *E. coli* showing that PG precursor composition can determine a cell’s preference for lateral vs septal cell wall growth (42-44). For instance, increasing the levels of a D,D-carboxypeptidase, which cleaves the terminal D-alanine on pentapeptide subunits to yield tetrapeptides, caused rod-shaped WT cells to grow as spheres (43), and allowed filamentous *pbp3* hypomorphs to shorten and divide more efficiently (42). These findings are consistent with the model that the Rod system prefers to act on PG with pentapeptide subunits and does not use tetrapeptides (or their tripeptide derivatives) efficiently (42, 43). The opposite may be true of the Divisome, in which the main transpeptidase PBP3 is thought to prefer or require tripeptides, originating from tetrapeptides, as the acceptor muropeptide (45, 46). Interestingly, dramatic Rod^-^ phenotypes are also seen with mutation of *A. baumannii dacC* (5), a predicted PG hydrolase that may also modulate the balance of cell wall peptide substrates in ways promoting lateral cell wall growth. The idea that cell wall subunit levels determine Rod system vs divisome activity may also contribute to the mitigation of toxicity in Δ*elsL* cells by Ldt_Ab_, in agreement with what was proposed in (6). While this mitigation is likely due to provision of alternative, supportive cross-links, an additional possibility is that Ldt_Ab_ directly or indirectly generates tripeptide substrates in Δ*elsL* that are needed for optimal cell division. Further work is required to understand the mechanisms by which alterations to cell wall stem peptides in *A. baumannii* drives the function (and malfunction) of specific PG biosynthesis enzymes.

Our genetic interaction analysis revealed an additional interdependent relationship between ElsL and the Mla lipid transport pathway. The Mla proteins mediate retrograde transport of phospholipids from the OM to the inner membrane, facilitating lipid asymmetry in the OM (23, 47). The system may also prevent excessive loss of lipids in the form of OM vesicles (OMVs) (48, 49). Interestingly, an *elsL* mutant sheds higher amounts of OMVs than WT (8). It is thus possible that membrane lipid loss contributes to the aggravating phenotype of the *elsL*/*mla* double mutant. Another possibility is that combining an impaired cell wall (due to low cross-linkage, ElsL^-^) with a compromised OM (Mla^-^) leads to synergistic failure of envelope mechanical integrity (50), manifesting as an aggravated growth defect. In sum, a variety of pathways, including those determining alternative crosslink formation, cell division, and OM homeostasis, become indispensable in the absence of ElsL function, underlining the importance of this key enzyme to multiple facets of *A. baumannii* envelope biogenesis.

The novel ElsL class of cell wall recycling LDC identified in this work uses a cysteine catalytic residue, contrasting it from that of the two described recycling LDC enzymes, LdcA and LdcV. LdcA, found in *E. coli, Pseudomonas aeruginosa*, and several other organisms, is a serine peptidase that uses a Ser-His-Glu catalytic triad (51), while LdcV, identified recently in *Vibrio cholerae, Aeromonas hydrophila* and *Proteus mirabilis*, belongs to the LAS superfamily of zinc-dependent metalloproteases that use an active site zinc ion coordinated by histidines (6, 52). While all three LDC classes enable muropeptide recycling, they may differ in their relative preferences for the variety of potential tetrapeptide-containing substrates derived from the cell wall (7), although this remains to be determined. The identification of a distinct, non-canonical class of LDC related to LDTs has two important implications. First, the finding helps illuminate the muropeptide recycling pathway in many bacteria, like *A. baumannii*, that lack a homolog of the previously identified recycling LDCs (33). Lack of a known LDC homolog in such bacteria has been explained previously by the possibility that they bypass the need for a recycling LDC due to low cell wall tetrapeptide content or high Mpl selectivity against tetrapeptides (33). While these mechanisms may be true in many cases, an alternative explanation is that a range of organisms possess a non-classical LDC (e.g., of the ElsL family), allowing the typical peptide recycling loop to be fully closed. Second, the reliance of *A. baumannii* on two YkuD-domain proteins for maintaining the cell wall points to a unique potential vulnerability in this highly resistant pathogen. Simultaneously targeting the catalytic cysteines in both ElsL and Ldt_Ab_ could exploit the synthetic lethality of the two proteins to potently weaken the sacculus of the pathogen and inhibit growth. Copper (14) and carbapenem antibiotics (53), known inhibitors of LDT proteins, are candidates for such a strategy.

In conclusion, we have identified a new family of LDC critical to muropeptide recycling in *A. baumannii* and likely many other organisms. Our combined genetic and biochemical interrogation of ElsL function provided insights into the role of the protein in recycling as well as its pleiotropic influence on numerous pathways important to envelope integrity and growth. Future work will examine the mechanisms by which changes to cell wall precursors in cells showing altered expression of ElsL or other muropeptide modifying enzymes lead to preferential action/inaction of specific PG synthesis machines, which will likely shed light on how cell wall synthesis and cell proliferation are coordinated in *A. baumannii*. Furthermore, the vulnerabilities exposed by ElsL inhibition, precursor buildup, and their network of toxic effects present opportunities for the rational development of improved antimicrobials aimed at controlling this intractable pathogen.

## Materials and Methods

### Bacterial strains and growth conditions

Bacterial strains used in this work are described in Table S2. *A. baumannii* strains were derivatives of ATCC 17978. Bacteria were cultured in Lysogeny Broth (10 g/L tryptone, 5 g/L yeast extract, 10 g/L NaCl) (LB), unless otherwise noted. Liquid cultures were incubated at 37°C in flasks with orbital shaking or in tubes with rotation via roller drum. Growth was monitored by measuring absorbance at 600nm. LB agar was supplemented with antibiotics (carbenicillin [Cb] at 25-100 μg/ml, kanamycin [Km] at 10-20 μg/ml, gentamicin [Gm] at 10 µg/ml) or sucrose (10%) as needed (Sigma Aldrich). LB prepared without NaCl (LB_0_) was used in phenotypic testing where noted. Super optimal broth with catabolite repression (SOC) was used in Tn-seq library preparation, and Vogel Bonner Medium (VBM) supplemented with Gm (10 µg/ml) was used for isolation of CRISPRi strains containing miniTn7 constructs.

### Molecular cloning and strain construction

Plasmids used in this study are listed in Table S2. Most DNA constructs were generated by PCR-amplification using oligonucleotide primers (Table S2) and cloning in the HincII site of pUC18, followed by verification by sequencing (Genewiz) before subcloning in subsequent vectors. Plasmids for complementation, localization, allele exchange, and sgRNA production were introduced into *A. baumannii* ATCC 17978 by electroporation (54).

#### Complementation and localization experiments

To generate an *ldt*_*Ab*_ plasmid for HADA experiments, the EcoRI and PstI fragment from pYDE231 was subcloned downstream of the T5lacP promoter in pEGE305 to generate pYDE240. To construct *elsL-* and *ldcA*-3xFLAG fusions for complementation experiments, the BamHI-XbaI fragment from pYDE342 and pYDE343 were each subcloned in pJE42, which provides an in-frame C-terminal 3xFLAG sequence, generating pYDE346 and pYDE347. The hybrid genes were then subcloned into pEGE305 using EcoRI and PstI, resulting in pYDE350 (ElsL_FLAG_) and pYDE351 (LdcA_FLAG_). To construct *msGFP2* gene fusions, a codon-optimized *msGFP2* reporter gene containing an in-frame poly-glycine linker was first synthesized as a double-stranded DNA fragment by IDT and cloned in the HincII site of pUC18 (pAFE225). The BamHI-XbaI fragments from pYDE342 and pYDE063 were cloned in pAFE225, generating pYDE386 (*elsL*-*msGFP2*) and pYDE387 (*ldt*_*Ab*_-*msGFP2*). The gene fusions were then cloned into pEGE305 using EcoRI and PstI, generating pYDE389 and pYDE390, respectively. To generate an independent *msGFP2* control gene, *msGFP2* was amplified from pAFE225 using a primer (msGFP2_F) that replaces the XbaI site and glycine linker with EcoRI, SD and translational start sites. The PCR product was then directly cloned into pEGE305 using EcoRI and PstI sites, generating pAFE256. An *elsL*_FLAG_(C138S) mutant was constructed by replacing the EcoRI-BstBI fragment of pYDE350 with a fragment amplified with a primer (elsL-C138S-R) containing the substitution mutation.

#### *Overexpression and purification of elsL* and *ldcA*

The NcoI-EcoRI fragments from pYDE281 and pYDE324, respectively, were each cloned upstream of the in-frame 6xHis tag sequence in pET28b, creating (pYDE290 and pYDE328).

#### Gene deletions

Deletion mutants were isolated using homologous recombination/allelic exchange using sucrose counterselection as described (55). A Δ*elsL* Δ*ldt*_*Ab*_ double mutant (EGA740) was constructed by allelic exchange with pEGE268 in EGA739 (5); the strain was isolated as small, sucrose-resistant colonies after overnight 30°C incubation followed by one day at room temperature. A Δ*elsL* Δ*ampG* double mutant (YDA411) was isolated by allelic exchange with pEGE268 in EGA516 (56). A Δ*elsL* Δ*ldt*_*Ab*_ Δ*ampG* triple mutant (YDA414) was isolated by allelic exchange with pEGE271 in YDA411. In-frame deletion of *ltgF* was constructed by three-way ligation of ∼1 kb flanking homology arms with pJB4648.

#### CRISPRi

sgRNAs were constructed by PCR-amplifying 24-base, PAM-adjacent target regions satisfying previously described criteria (57, 58), followed by cloning directly into the SpeI and ApaI sites of pYDE007 (25) and verification by restriction digestion and sequencing. Δ*elsL* and Δ*ldt*_*Ab*_ strains with inducible *dcas9* were isolated by introducing pYDE009 (25) into EGA738 and EGA739 via four-parental mating (55, 59); transposants were isolated on VBM-Gm10 plates, generating YDA186 and YDA095, respectively. Integration at the *attTn7* locus was confirmed by PCR and loss of pYDE009 was confirmed by screening on Cb plates (60, 61).

### Construction of transposon mutant libraries

Mutagenesis of Δ*elsL* (EGA738) and Δ*ldt*_*Ab*_ (EGA739) with the *mariner* transposon was performed by electroporation with pDL1100 (5). Cells were then diluted with 1 ml SOC, allowed to incubate for 15 min at 37°C, and spread on membrane filters (0.45 μm pore size) overlaid on prewarmed SOC agar plates. Plates were incubated at 37°C for 1 h, and filters were transferred to solid LB supplemented with Km (10 µg/ml with Δ*elsL*, and 20 µg/ml with Δ*ldt*_*Ab*_). After overnight incubation at 37°C, colonies were lifted from filters by agitation in sterile PBS, mixed with sterile glycerol (10%), aliquoted, and stored at -80°C. Approximately 160,000 mutant colonies from 19 subpools were analyzed in the Δ*elsL* strain, and approximately 300,000 from 20 subpools were analyzed with the Δ*ldt*_*Ab*_ strain.

### Tn-seq library amplification, sequencing, and analysis

Genomic DNA was extracted from samples using the DNeasy Kit (Qiagen) and quantified by a SYBR green I (Invitrogen) microplate assay. Transposon-adjacent DNA was tagmented and amplified for Illumina sequencing as described (5, 62). Samples were multiplexed, reconditioned, and size selected (250- or 275-600bp, Pippin HT) before sequencing (single-end 50bp) using primer mar512 on a HiSeq2500 with High Output V4 chemistry at Tufts University Genomics Core Facility.

Sequencing reads were quality-filtered, clipped of adapters, and mapped to the *A. baumannii* chromosome (NZ_CP012004) with Bowtie (25). Mapped reads were tabulated in wig format according to the position of their TA sites in the NZ_CP012004 genome using custom python scripts (25). To examine genetic interactions between transposon mutations and the deleted gene, datasets were analyzed by the Resampling method in the TRANSIT software package (parameters: samples=10000000, norm=TTR, histograms=False, adaptive=True, excludeZeros=False, pseudocounts=0.0, LOESS=False, trim_Nterm=0.0, trim_Cterm=10.0) (63), using previously published WT *mariner* libraries as the control dataset (25). Resampling p values were adjusted for multiple comparisons (q value) using the method of Benjamini and Hochberg. Results were visualized as volcano plots using Prism 8. Genes having low mean read count (< 5) in both the deletion and control datasets (essential genes) were not plotted. Hits were defined as genes having q value < 0.05; ≥ 3 transposon sites; and > 5 mean read counts in both datasets. To visualize Tn-seq read counts along chromosomal regions, TTR-normalized counts were merged into a single wig file, scaled such that median read coverage at non-zero insertion sites was equivalent between datasets, and viewed using Integrative Genomics Viewer (25, 64).

### Microscopy

Bacteria in the log phase of growth were immobilized on agarose pads (1% in PBS) prior to imaging. For HADA labelling experiments, bacteria were pulsed with 1 mM HADA (Tocris Bioscience) for 15 minutes, fixed with 70% ice-cold ethanol for 10-15 minutes, washed twice with PBS before imaging. For localization experiments, strains harboring IPTG-inducible gene fusions to msGFP2 were cultured with 100 µM IPTG prior to imaging. Micrographs were acquired with a 100x/1.4 phase-contrast objective on a Zeiss Axio Observer 7 microscope. A Colibri 7 LED light source was used for fluorescence illumination. Imaging of GFP fluorescence used the 475 nm LED and filter set 92 HE. Imaging of HADA fluorescence used the 385 nm LED and filter set 96 HE. For analysis of HADA incorporation, Fiji software (65) was used for background subtraction from the HADA signal and for determining cell boundaries from stacked HADA and phase images. Fluorescence intensity across populations of cells in multiple fields was then plotted as demographs using MicrobeJ software (66). For analysis of cell width, the maximal width relative to the medial axis of each cell was quantified from phase images using MicrobeJ (66).

### Genetic interaction and antibiotic susceptibility tests

Bacteria (OD 1) were serially diluted 10-fold in PBS and spot-plated on the noted solid medium. After overnight growth at 37°C, colonies were imaged with transilluminated light on a ChemiDoc MP Imaging System (BioRad). For genetic interaction analysis, colony diameters were measured using ImageJ (67) and normalized to the WT control (YDA007). Genetic interactions were analyzed by comparing the relative colony diameter values of strains harboring two genetic lesions (CRISPRi knockdown of candidate gene and deletion of Δ*elsL*) with the hypothetical values expected from a multiplicative model, in which the values of each single-lesion strain are multiplied (68). Sensitivity to sulbactam and to piperacillin-tazobactam (8:1 ratio by mass) was measured by CFE assay (56). Serial dilutions of WT and isogenic mutant strains were spotted on solid LB agar medium containing graded concentrations of drug (or no drug control). After overnight growth at 37°C colony counts were determined and compared to no drug control. Limit of detection was approximately 10^−5^ to 10^−6^.

### PG isolation and analysis

Sacculi isolation and PG analysis was performed as previously described (69). In short, *A. baumannii* cells from overnight LB cultures were harvested, resuspended in LB + 5% SDS, and boiled with stirring for 2 hours followed by stirring at room temperature overnight. SDS was removed from sacculi through several rounds of ultracentrifugation and resuspension with MilliQ water. Sacculi were then resuspended in 100 mM Tris-HCl pH 8.0 buffer with proteinase K (20 µg/ml) and incubated at 37°C for one hour. The reaction was stopped by adding SDS 1% and boiling at 100°C for 5 minutes. SDS was removed as above and sacculi were resuspended in water. Muramidase was then added and reactions incubated overnight at 37°C to solubilize the sacculi completely. The soluble muropeptides were reduced using NaBH_4_ and their pH adjusted before separation using UPLC (Waters) and identification using a MALDI-TOF MS system (Waters).

For quantification, we chose 2 random PG profiles that were representative of each strain. The area for each identified peak was integrated, giving each individual muropeptide a relative area value based on the total integrated area. Using these values, the molar percentage was also calculated for each muropeptide. This relative molarity was also used to calculate the degree of crosslinking using the formula: 

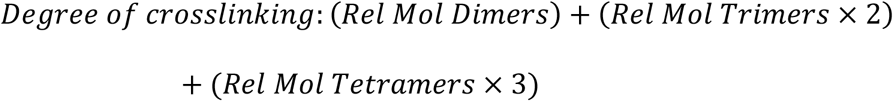

### Analysis of intracellular and extracellular soluble muropeptides

Bacteria were grown with LB at 37°C for 4 hours to log phase and OD recorded. Bacteria were cooled on ice and centrifuged at 25,000 rpm for 20 minutes at 4°C. Supernatants were stored at 4°C for analysis of extracellular soluble muropeptides, and pellets were used for analysis of the intracellular fraction.

To study PG turnover products and precursors within the intracellular fraction, cell pellets were washed twice with sterile 1% NaCl, resuspended in water, and boiled for 30 minutes with stirring. Samples were again centrifuged at 25,000 rpm for 15 minutes and supernatants were recovered and filtered (0.2 µm pore-size). A MALDI-TOF MS system was used for identification of the muropeptides derived from turnover (anhydro species) and synthesis (UDP-activated species). To study muropeptides within the extracellular fraction, supernatants were filtered as above and immediately boiled for 15 minutes to precipitate proteins. The soluble fraction was then analyzed using the same MS system as for the intracellular muropeptides, focusing on detection of products derived from turnover that were released into the media. Quantification of all muropeptides used the total-ion count detected by the system. All analyses were performed with biological triplicate samples.

### Suppressor mapping by whole-genome sequencing and phenotypic signature analysis

Suppressor mutants bypassing the EGA740 growth defect were isolated on LB or LB_0_ plates. Genomic DNA (DNeasy Kit) was quantified by a SYBR green I (Invitrogen) microplate assay, and used as input for Illumina library preparation using a modified small-volume Nextera tagmentation method as described (56). Libraries were sequenced (single-end 100bp) on a HiSeq2500 at TUCF-Genomics. Reads were aligned to the NZ_CP012004 genome and variants identified using BRESEQ (70). The predicted functional impact of substitution variants was determined by using PROVEAN (71). Identities of IS sequences was determined by using ISFinder (72). Phenotypic signatures of *ltgF* and cell wall recycling genes were compared using Qlucore Omics Explorer (3.5), and Pearson correlations were analyzed with Prism 8.

### Immunoanalysis of ElsL, Ldt_Ab_, and LdcA proteins

Strains harboring fusion genes or vector control were diluted to OD 0.05 in LB with IPTG (100 µM with msGFP2 fusions; 0, 50, or 500 µM with 3xFLAG fusions) and grown to OD 0.5. Cells were centrifuged and resuspended (50 µl per 1 mL of sample) with SDS sample loading buffer (msGFP2 fusions) or BugBuster Protein Extraction Reagent with 0.1% lysonase (Millipore) followed by SDS loading buffer (3xFLAG fusions). Samples were boiled for 10 minutes, separated by SDS-PAGE (12% acrylamide gel), and transferred to an Immobilon-FL PVDF membrane. Total protein was detected by SYPRO Ruby Protein Blot Stain (Invitrogen). GFP was detected by rabbit PABG1 primary (Chromotek, 1:5000 dilution) and goat anti-rabbit IgG HRP secondary (Invitrogen, 1:5000 dilution) antibodies. FLAG epitope was detected by mouse anti-FLAG M2 primary (Invitrogen, 1:1000 dilution) and goat anti-mouse IgG HRP secondary (Invitrogen, 1:5000 dilution) antibodies. Blots were imaged with a ChemiDoc MP (BioRad). Band intensities were quantified using Image Lab software. Samples were normalized by dividing the immunodetected band intensity by the total protein level from SYPRO staining. Relative values were calculated by dividing each normalized value by the normalized value of the ElsL_FLAG_, 50 µM IPTG sample on the same blot.

### Protein overexpression and purification

Overnight cultures of *E. coli* BL21(DE3) strains harboring pYDE290 (*elsL*_*His*_) or pYDE328 (*ldcA*_*His*_) were seeded at 1:100 dilution into 1L of LB with Km (30 µg/mL) and grown at 37°C to OD 0.6-0.8. IPTG was added at 500 µM final and cultures were incubated at 16°C for 20 hours. Cells were harvested, resuspended with 30 mL of pre-chilled buffer A (50 mM Tris-HCl, 300 mM NaCl), and pulse sonicated on ice for 15 minutes. The lysate was clarified by centrifugation at 10000 g at 4°C for 1 hour and filtered (0.45 µm pore size, Millipore). Clarified lysates were loaded on Ni-NTA resin columns (Thermo Scientific) previously equilibrated with 6% (v/v) Buffer B (50 mM Tris-HCl, 300 mM NaCl, 500 mM imidazole) in Buffer A. His-tagged proteins were eluted with gradient concentrations of Buffer B (6%, 10%, 30%, 50%, 100%) at 4°C, and detected by SDS-PAGE and Coomassie Blue staining. Fractions containing the His-tagged proteins were concentrated by using Amicon Ultra-15 Centrifugal Filter Units (10 kD). Proteins were washed 3 times with 15 mL stocking buffer (50 mM Tris-HCl, 50 NaCl, 10% (w/v) glycerol), resuspended with 200 µL stocking buffer. Proteins were quantified using the Pierce Coomassie Bradford Protein Assay (Thermo Scientific) and stored as aliquots at -80°C. With ElsL_His_, the elution and stocking buffers included 0.5 mM tris(2-carboxyethyl)phosphine (TCEP) to prevent oxidation of the active site cysteine residue (73).

### *In vitro* assays with ElsL

LDC activity was assayed *in vitro* using purified proteins and muropeptide substrates. Muropeptide substrates (M4 and M5) were extracted from sacculi isolated from *Caulobacter crescentus*, which are known to have high M4 and M5 content (74), and solubilized as described above. Soluble muropeptides were separated using a High Performance Liquid Chromatography (Waters) system, and each individual muropeptide was collected manually. Collected fractions were washed from solvents by evaporation using a SpeedVac vacuum concentrator followed by resuspension with MilliQ water. All reactions were performed in 100mM Tris-HCl, mixing a fraction of M4 or M5 and 10 ng of the corresponding protein (ElsL_His_, LdcA_His_ or none in the negative control). Reactions were incubated for 2 hours at 37°C and stopped by incubation at 100°C for 5 minutes. After inactivation, the samples were reduced and their pH adjusted for injection on the UPLC. Muropeptides were identified according to their retention time, and their identities were validated through MS analysis.

### ElsL ortholog identification

*A. baumannii* ElsL (WP_000077301) was queried by BLASTp using the refseq_select_prot database, restricting the search to Bacteria (representing ∼14K genomes) and an *E*-value cutoff of 1e-4. From the 1082 protein sequences that were returned, custom python scripts were used to identify sequences containing a predicted signal peptide or transmembrane helix (based on the SignalP-5.0 (75), PrediSi (76), Phobius (77), and TMHMM 2.0 (78) methods), or PG-binding domain (from search of the NCBI Conserved Domain Database (CDD) (40)). With SignalP-5.0, an “other” score of < 0.75 was used as a conservative marker of potential signal peptides. Hits were excluded if any of the above searches were positive. Finally, sequences shorter than 105 or longer than 200 amino acids were excluded. Taxonomic information for the resulting set of ElsL orthologs (788 hits) was obtained from the NCBI database.

## Data Availability

All sequence data can be found in the NCBI Sequence Read Archive under accession code PRJNA763919.

## Acknowledgements

This work was supported by Northeastern University startup funds and by the NIAID/NIH under award number R01AI162996 to E.G. Research in the Cava lab is supported by the Swedish Research Council, The Laboratory of Molecular Infection Medicine Sweden (MIMS), The Knut and Alice Wallenberg Foundation (KAW), Umeå University and the Kempe Foundation. We thank Sylvie Manuse and Emily Kirkwood for assistance with microscopy and image analysis, Wei-Leung Ng for plasmid gifts, Tom Bernhardt and David Roper for experimental advice, and members of the Geisinger lab for helpful discussions.

## Supporting Information

**Fig. S1. UPLC/MS analysis of *A. baumannii* cell wall**. Muropeptide fragments isolated from Δ*elsL* sacculi were identified by MS, and the corresponding peaks in the UPLC chromatogram (bottom; same as that in Fig. 1E) were labelled accordingly (blue numbers). The reference WT UPLC chromatogram is also shown (top; same as that in Fig. 1E).

**Fig. S2. Western blot analysis of fusion protein levels**. WT or Δ*elsL* bacteria containing the indicated construct driven by P(IPTG) were cultured with the noted IPTG dose, and proteins were analyzed after blotting. The top image in each panel shows representative immunoblot using indicted primary antibody. Bottom image shows total protein levels from the same blot determined by SYPRO Ruby staining prior to immunodetection. Location of MW markers (BioRad Precision Plus Protein Dual Color) are indicated to the left of each blot. -, no insert (vector control). NA, not analyzed. (A) Detection of msGFP2 fusion proteins. (B-C) Detection of 3xFLAG epitope-tagged proteins. Arrowheads indicate position of the indicated fusion protein (ElsL_FLAG_ in C).

**Fig. S3. Analysis of *elsL* genetic interactions with *RS13190-zapA* and *ldt***_***Ab***_. (A) *elsL* deficiency blocks elongated phenotype caused by *RS13190-zapA* knockdown. WT or Δ*elsL* bacteria harboring *dcas9* and the indicated sgRNA were cultured in the presence of aTc (50 ng/mL) and imaged in log phase by phase-contrast microscopy. Scale bar, 4 µm. (B) *elsL* and *ldt*_*Ab*_ are synthetic lethal on LB_0_ medium. Genetic interaction was analyzed by CRISPRi knockdown of *ldt*_*Ab*_ in the Δ*elsL* strain background. Colonies were grown on LB_0_ agar plates and imaged as in Fig 3C. (C) Muropeptide profile analysis of the Δ*elsL* Δ*ldt*_*Ab*_ strain compared to single mutants and WT analyzed in parallel. The muropeptide profiles of the single mutants and WT are the same as those shown in Fig. 1E and are included for comparison with Δ*elsL* Δ*ldt*_*Ab*_. Muropeptides are labelled as in Fig. 1E. (D) Percentage of crosslinked muropeptides determined from muropeptide profiles (Materials and methods). First 3 samples are the same as those shown in Fig. 1H and are included for comparison with Δ*elsL* Δ*ldt*_*Ab*_. Bars show mean ± s.d.

**Fig. S4. Phenotypes of LtgF (ACX60_RS13100) are closely related to those of cell wall recycling genes**. (A) Tn-seq phenotypic signature of *ltgF* clusters with those of cell wall recycling genes. Each row is the Tn-seq phenotypic signature of the indicated gene. Heat map shows normalized Tn-seq fitness in z-scored units across a panel of distinct drug treatments (columns). Phenotypic signature data are from (5). (B-C) Disruption of *ltgF* in strain AB5075 results in hypersusceptibility to fosfomycin. (B) AB5075 WT or indicated mutant were serially diluted and spotted on solid medium without or with drug, and resulting colonies were imaged as in Fig. 3. (C) Growth of bacteria in panel B was monitored during culture in microplates with or without fosfomycin (75 µg/ml). Data points show geometric mean ± s.d. (dotted bands) from n = 2 independent cultures. Where not visible, s.d. is within the confines of the symbol. (D) *ldt*_*Ab*_ genetic interactions with cell wall recycling genes are not dependent on the component being recycled. Heat map shows fold change in Tn-seq read counts of the indicated gene in the Δ*ldt*_*Ab*_ *mariner* library vs the WT control library.

**Fig. S5. Responses of *A. baumannii* WT and mutants to piperacillin-tazobactam indicates that the treatment targets divisional cell wall synthesis**. (A) Piperacillin-tazobactam, like sulbactam, causes WT cells to grow as filaments, consistent with targeting divisome PG synthesis. Cells were grown with or without sub-MIC piperacillin-tazobactam and were imaged by phase-contrast microscopy. Scale bar, 10 µm. (B-C) ElsL and PBP2 mutants show similar hypersusceptibility to piperacillin-tazobactam. Susceptibility was measured by CFE assay. Data points show geometric mean ± s.d. (n = 3 biological replicates).

**Table S1. Quantification of genetic interactions with Δ*elsL*.**

**Table S2. Strains, plasmids, and primers used in this study.**

**Dataset S1. Tn-seq genetic interactions with Δ*elsL***.

**Dataset S2. Tn-seq genetic interactions with Δ*ldt***_***Ab***_.

**Dataset S3. ElsL orthologs**.

